# Ubiquitin ligase CHFR impairs Tie2 signaling via K^48^-linked ubiquitylation and degradation of Akt1 in endothelial cells

**DOI:** 10.64898/2026.03.31.715582

**Authors:** Mohammad Owais Ansari, Gary C.H. Mo, Lasanthi Jayathilaka, Hui Chen, Asrar B. Malik, Chinnaswamy Tiruppathi

## Abstract

Vascular endothelial (VE)-cadherin is essential for maintaining endothelial junctional barrier integrity. The Angiopoietin-1 (Ang-1)/Tie2 axis induced Akt1 activation is crucial for maintaining endothelial junctional barrier by inhibiting FoxO1 and suppressing expression of Angiopoietin-2 (Ang-2), a Tie2 antagonist. Systemic inflammatory conditions such as sepsis, Akt1 expression is reduced, whereas FoxO1-dependent Ang-2 expression is increased, resulting in endothelial barrier dysfunction. We previously showed that the TLR4/FoxO1 axis induces the ubiquitin E3 ligase CHFR, which promotes endothelial barrier disruption by targeting VE-cadherin for ubiquitylation and degradation. However, little is known about Akt1 expression during vascular inflammation. Here, we identified FoxO1-dependent CHFR expression as a key mechanism driving K48-linked polyubiquitylation and proteasomal degradation of Akt1 in endothelial cells (EC). LPS-induced K^48^-linked ubiquitylation of Akt1 was prevented in CHFR-depleted human EC and in endothelial-specific *Chfr* knockout (*Chfr^ΔEC^*) mice. Accordingly, CHFR depletion increased Akt1 and VE-cadherin expression in both human lung EC and *Chfr^ΔEC^* mice. *Chfr^ΔEC^* mouse lungs also exhibited elevated Ang-1 and Tie2 expression, and Ang-1 stimulation induced sustained Akt1 phosphorylation in CHFR-deficient EC. Moreover, CHFR depletion prevented LPS-induced expression of FoxO1 and Ang-2 in EC. Mechanistically, CHFR interacted with phosphorylated Akt1 and mediated its ubiquitylation at lysine residues K30, K39, K154, and K268. Expression of a ubiquitylation-deficient Akt1 mutant prevented LPS-induced VE-cadherin degradation and vascular injury. Collectively, these findings identify CHFR as a critical regulator of endothelial inflammatory responses by controlling Akt1 stability and VE-cadherin expression during inflammation.

## Introduction

The serine/threonine kinase Akt (also known as protein kinase B, PKB) plays a central role in endothelial cell (EC) biology (*1*). Mammals express three closely related isoforms Akt1 (PKBα), Akt2 (PKBβ), and Akt3 (PKBγ) encoded by distinct genes but sharing ∼80% sequence identity and similar structural organization (*2*). Several studies confirm Akt1 is the major isoform in EC from different vascular beds (*3*). Akt2 is also expressed in EC, but typically at lower levels than Akt1. Akt3 is much less expressed or nearly negligible in EC of many organs. Importantly, reduced postnatal and ischemia-induced angiogenesis, EC apoptosis, and impaired migration were observed in Akt1 knockout mice (*4*). Further, in EC-specific Akt1 knockout mice, impaired eNOS activation and endothelial barrier dysfunction was observed (*4*). Furthermore, phospho-proteomic studies showed that in EC, Akt1 preferentially phosphorylates several substrates relevant for angiogenesis, eNOS, and transcription factor FoxO1 (*1*, *5*). Collectively, these findings underscore the fundamental role of Akt1 in regulating endothelial functions.

Vascular homeostasis requires tight control of EC survival, barrier function, and response to stress. Among endothelial receptor systems, the angiopoietin-Tie2 signaling axis uniquely maintains vascular barrier stability (*6*). Tie2 (also known as Tek) is an endothelium-specific receptor tyrosine kinase whose ligands, Angiopoietin-1 (Ang-1) and Angiopoietin-2 (Ang-2), function as context-dependent agonist and antagonist, respectively (*7*). The dynamic balance between Ang-1/Tie2 activation and Ang-2-mediated inhibition of Tie2 determines whether vessels remain quiescent or become permeable and angiogenic (*8*). Tie2 prevents systemic inflammation-induced thrombus formation in the lung vasculature (*9*). Tie2 expression at the endothelial surface is required to maintain vascular endothelial cadherin (VE-cadherin, a.k.a. CDH5) expression at endothelial adherens junctions (AJs) (*10*, *11*). The loss of VE-cadherin at AJs results in endothelial barrier breakdown, leading to uncontrolled leakage of plasma proteins into tissue and the influx of blood cells (*12*, *13*). Interestingly, *in vivo* activation of Tie2 with recombinant Ang-1 or lipid nanoparticle-mediated *in vivo* delivery of mRNA encoding Ang-1 into lung vascular endothelium prevented lung vascular leak, pulmonary edema, and neutrophil infiltration induced by LPS challenge in mice (*14*, *15*). Further, studies have shown that Tie2 expression and function rapidly declined in many models of vascular leak-associated infections including influenza, SARS CoV2 (COVID-19), and sepsis (*16*, *17*). However, the molecular basis of downregulation of Tie2 signaling pathways in EC during vascular inflammation remain poorly understood.

Pericytes are specialized cells, in close contact with EC within the basement membrane of microvasculature is crucial for endothelial barrier stability (*18–20*). The release of Angiopoetin-1 (Ang-1) from pericytes constitutively activates endothelial enriched receptor tyrosine kinase Tie2 and its downstream PI3K-Akt1 axis, which in turn promotes endothelial barrier stability through the expression of VE-cadherin at endothelial AJs (*21–23*). Akt1-mediated phosphorylation of the transcription factor FoxO1 prevents transcriptional function by promoting nuclear exclusion, ubiquitylation, and proteasomal degradation, which increases the expression of genes that cause vascular stabilization (*24*, *25*). In addition, Akt1 inactivates GSK3β through phosphorylation of GSK3β on Ser-9 to block β-catenin phosphorylation and degradation via proteosome which in turn strength the interaction between β-catenin and VE-cadherin at EC AJs (*26*, *27*). These findings indicate that endothelial Akt1 signaling plays a vital role in vascular barrier stability. Importantly, systemic inflammatory conditions such as sepsis, Akt1 expression and function were remarkably suppressed while FoxO1-mediated increased expression of Tie2 antagonist Ang-2 in EC disrupts endothelial barrier integrity (*7*, *25*). However, little is known about the biochemical mechanisms that promote the dysregulation of endothelial barrier protective Akt1 expression and function in EC during vascular inflammation.

The ubiquitin-proteasome system (UPS) plays a central role in regulating protein turnover to maintain cellular homeostasis and proper cellular function. The UPS is composed of three main components: E1 (Ubiquitin-activating enzyme) initiates the process by activating ubiquitin; E2 (Ubiquitin-conjugating enzyme) transfers ubiquitin from E1 to the target protein; and E3 ligase (Ubiquitin-protein ligase) responsible for the specific recognition and transfer of ubiquitin from E2 to the target protein. Ubiquitin E3 ligase TRAF6 mediated ubiquitylation of Akt1 via K^63^-linked monoubiquitylation was shown to activate Akt1 (*28*). In contrast, in cancer cell lines and fibroblast, ubiquitylation of Akt1 via K^48^-linked polyubiquitylation by E3 ligases such as MULAN, TTC3, and TRIM36 induced degradation of Akt1 via proteasome (*29–31*). We have recently shown that the transcription factor FoxO1 mediated upregulation of ubiquitin E3 ligase CHFR during vascular inflammation induced ubiquitylation-dependent degradation of VE-cadherin in EC to cause endothelial barrier breakdown (*13*). Importantly, we showed that in EC-restricted *Chfr* knockout (*Chfr^τ<EC^*) mice, augmented expression of VE-cadherin and abrogation of LPS-induced degradation of VE-cadherin (*13*). Since VE-cadherin itself is known to positively regulate Akt1 function in EC (*32*, *33*), in this study, we investigated whether the EC-expressed CHFR also modulates the expression and function of Akt1 in EC and its consequences on endothelial barrier stability. Herein, we identified TLR4-FoxO1 axis induced CHFR expression is causing K^48^-linked ubiquitylation and degradation of Akt1 in EC. Further, we found that CHFR interacts with phosphorylated Akt1 and facilitates its ubiquitylation at lysine residues K30, K39, K154, and K268. We also found that in both CHFR-depleted EC and *Chfr^ΔEC^* mice, LPS-induced expression of FoxO1 and its target gene Ang-2 was blocked. These results support a novel mechanism whereby CHFR stabilizes FoxO1 by promoting Akt1 degradation in EC. Collectively, our findings reveal that endothelial CHFR regulates vascular inflammatory responses by modulating the expression and stability of both VE-cadherin and Akt1, highlighting CHFR as a critical modulator of endothelial function during inflammation.

## Results

### Endothelial cell–expressed ubiquitin ligase CHFR promotes Ang-2 expression via suppression of Akt1 function

In endothelial cell-restricted *Chfr* knockout (*Chfr^ΔEC^*) mice, we observed augmented VE-cadherin expression and complete abrogation of LPS-induced VE-cadherin degradation (*13*). Because Akt1 expression and activity are critical for endothelial barrier stability, we investigated whether CHFR also regulates Akt1 expression to promote endothelial barrier disruption in response to LPS. We observed increased Akt1 expression accompanied by suppression of FoxO1 in *Chfr^ΔEC^* mice (**Fig. 1a, b**). Notably, LPS-induced Akt1 degradation was prevented in *Chfr^ΔEC^* mice (**Fig. 1c; Supplemental Fig. 1a**). Consistent with this, LPS-induced Ang-2 mRNA and protein expression was blocked in *Chfr^ΔEC^*mice (**Fig. 1d, e**). Furthermore, LPS-induced downregulation of Tie2 was also reduced in *Chfr^ΔEC^* mice (**Fig. 1e**).

**Figure 1.**
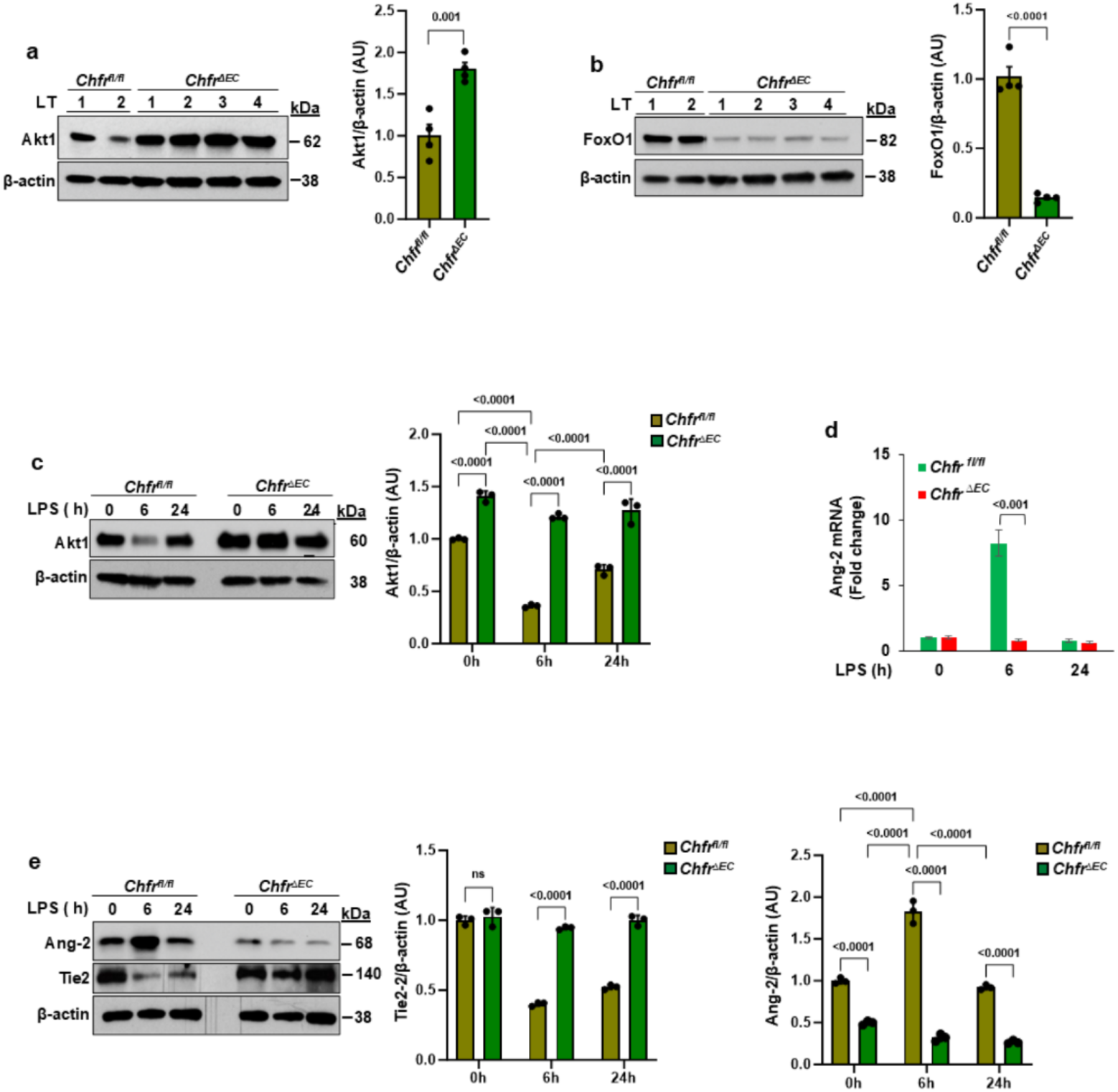
EC-restricted *Chfr* deletion in mice augments Akt1 expression and prevents expression of FoxO1 and Ang-2. **a-b** Immunoblot (IB) analysis of lung tissues (LT) showed increased expression of Akt1 and decreased expression of FoxO1 in *Chfr^ΔEC^*as compared to *Chfr^fl/fl^* (WT) mice (*n*=4 mice/genotype). Shown are mean values ± SEM (unpaired two-tailed Student’s t test). **C** *Chfr^fl/fl^* and *Chfr^ΔEC^* mice injected with LPS (10 mg/kg bodyweight, i.p.) for 0, 6, and 24 h were used to measure the expression of Akt1 in lung tissue. Shown are mean values ± SEM (*n* = 3 mice/genotype/group; two-way ANOVA followed by Tukey’s post-hoc test). **d-e** *Chfr^fl/fl^* and *Chfr^ΔEC^* mice were i.p. injection of LPS (10 mg/kg bodyweight) for 0, 6, and 24 h. Thereafter, lung tissues harvested were used for RT-qPCR to measure the mRNA expression of Ang-2 (**d**) or IB analysis was performed to determine the expression of Ang-2 and Tie-2 (**e**). Shown are mean values ± SEM (*n* = 3 mice/genotype/group; two-way ANOVA followed by Tukey’s post-hoc test).

Since VE-cadherin expression at endothelial AJs promotes pericyte recruitment to stabilize endothelial barrier integrity (*34*), we examined expression of the pericyte marker PDGFRβ in lungs of WT and *Chfr^ΔEC^* mice. LPS challenge induced pericyte loss in lung vessels of WT mice, whereas pericyte loss was not observed in *Chfr^ΔEC^* mice (**Supplemental Fig. 1b**). In addition, Ang-1 and Tie2 mRNA expression was significantly increased in lungs of *Chfr^ΔEC^* mice compared with WT controls (**Supplemental Fig. 1c**). Collectively, these findings suggest that CHFR suppresses Akt1 expression in response to LPS, thereby impairing Tie2 signaling and endothelial barrier integrity.

### Depletion of CHFR in endothelial cells augments Akt1 signaling

To determine whether CHFR depletion-mediated increases in Akt1 expression correlate with enhanced Akt1 signaling, we assessed Ang-1-induced Akt1 phosphorylation at Ser-473 and Thr-308 in control and CHFR knockdown human lung microvascular endothelial cells (HLMVEC). CHFR knockdown resulted in robust and sustained phosphorylation of Akt1 at both residues compared with control cells (**Fig. 2a, b**). Next, we measured live cell Akt activity utilizing Akt biosensor (*35*). Here, we observed basal as well as Ang-1 stimulated Akt activity was significantly increased in CHFR depleted HLMVEC compared with control HLMVEC (**Fig. 2c**). Consistent with enhanced signaling, Ang-1-induced Tie2 phosphorylation was also sustained in CHFR-depleted HLMVEC (**Fig. 2d**). Akt1-mediated phosphorylation of GSK3β at Ser9 inactivates GSK3β, thereby preventing β-catenin phosphorylation and proteasomal degradation, which stabilizes endothelial AJs (*27*, *36*). Accordingly, Ang-1-induced phosphorylation of GSK3β at Ser9 was markedly increased in CHFR-depleted cells compared with controls (**Fig. 2e**). These results support the conclusion that CHFR depletion enhances Akt1 signaling and strengthens endothelial AJ integrity.

**Figure 2.**
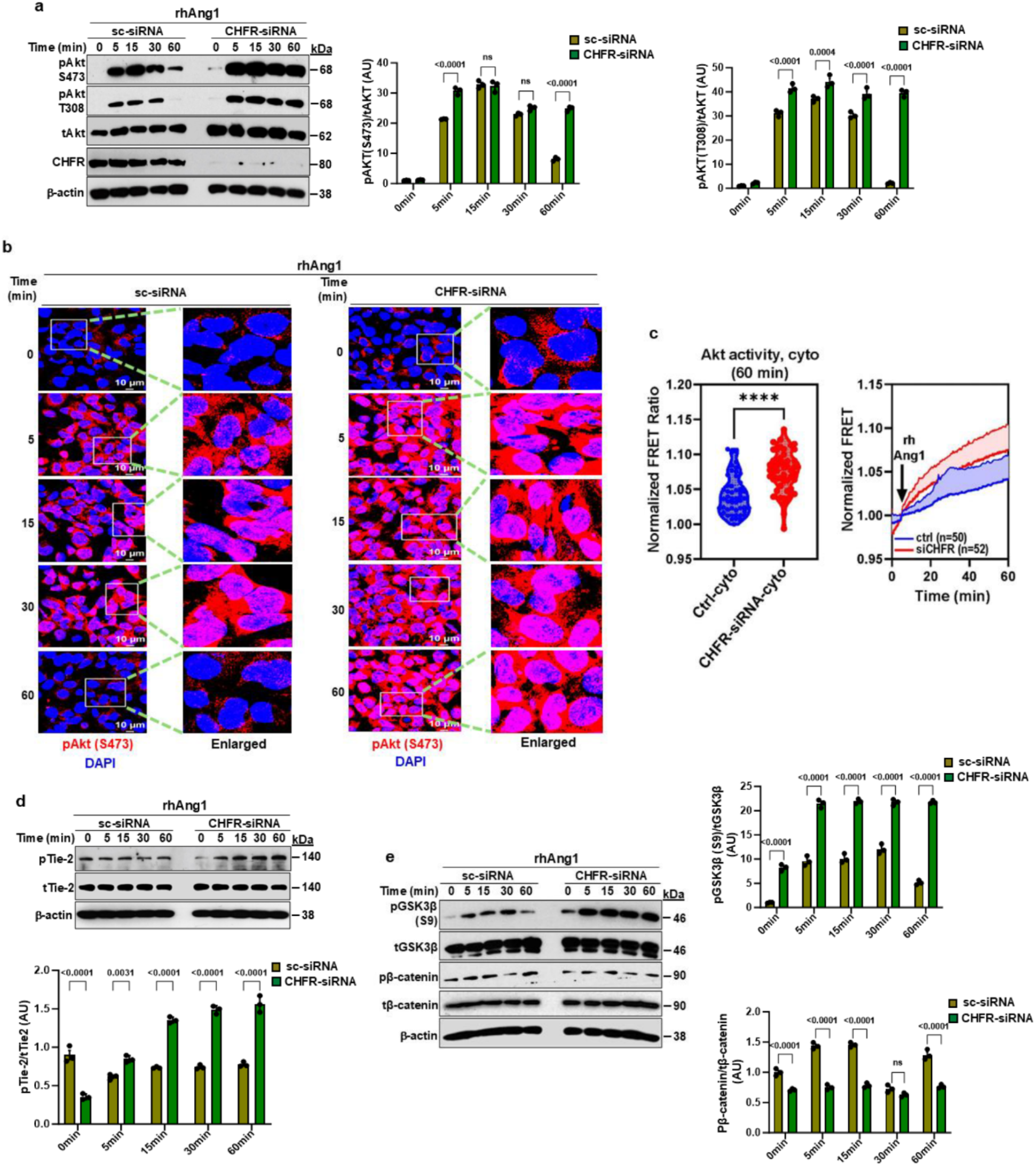
CHFR depletion augments Akt1 expression and function in endothelial cells. **a** HMEC were transfected with scrambled-siRNA (sc-siRNA) or CHFR-siRNA. At 72 h post-transfection, cells stimulated with rAng1 (400 ng/ml) for different time intervals were used for IB analysis to determine phosphorylation of Akt1 at S473 and T308 (*n* = 3 independent experiments). **b** Control HMEC and CHFR knockdown HMEC stimulated with rAng1 as above were stained with phospho-473-Akt1 specific antibody. **c** Control HMEC and CHFR knockdown HMEC were transfected with the Akt1 biosensor plasmid and then stimulated with rAng1 as above to assess live cell Akt1 activity by measuring the FRET ratio (*p*<0.0001) (*left panel* shows basal Akt activity; *right panel* shows Akt activity in response to rAng1 challenge). **d-f** Control HMEC and CHFR-siRNA treated HMEC stimulated with rAng1 were used for IB analysis to determine tyrosine phosphorylation Tie2 (**d**), phosphorylation of GSK3β at S9 and phosphorylation of β-catenin (**e**), (*n* = 3 independent experiments). **a, d-e** Shown are mean values ± SEM (*n* = 3 independent experiments; two-way ANOVA followed by Tukey’s post-hoc test).

### CHFR stabilizes FoxO1 by downregulating Akt1 in endothelial cells

We previously demonstrated that LPS-induced FoxO1 expression is required for CHFR induction in endothelial cells (*13*). FoxO1 is also essential for expression of the Tie2 antagonist Ang-2 during vascular inflammation in endothelial cells (*13*, *25*). Although our data show that LPS-induced Ang-2 expression is blocked in *Chfr^ΔEC^*mice (**Fig. 1**), the mechanism by which FoxO1-mediated Ang-2 expression is suppressed in the absence of CHFR remained unclear. Akt1 is known to negatively regulate FoxO1 through phosphorylation of FoxO1 in endothelial cells (*37*). Akt1 translocation to the nucleus, induces phosphorylation of FoxO1 at S256 inhibits FoxO1 binding to DNA whereas phosphorylation at T24 expels FoxO1 from nucleus binding to 14-3-3 protein and subsequently FoxO1 is degraded through ubiquitylation by E3 ligase skp2 (*37*). Thus, we examined whether CHFR-mediated Akt1 degradation contributes to FoxO1 expression and function. In control HLMVEC, LPS exposure caused marked downregulation of Akt1 at 6 and 12 h, whereas Akt1 downregulation was prevented in CHFR-depleted cells (**Fig. 3a**). Immunofluorescence analysis revealed concomitant loss of Akt1 and VE-cadherin in control cells at 6 and 12 h following LPS challenge, with recovery by 24 h (**Fig. 3b**). In contrast, robust nuclear localized Akt1 and VE-cadherin expression at AJs remained intact in CHFR-depleted HLMVEC (**Fig. 3b**).

**Figure 3.**
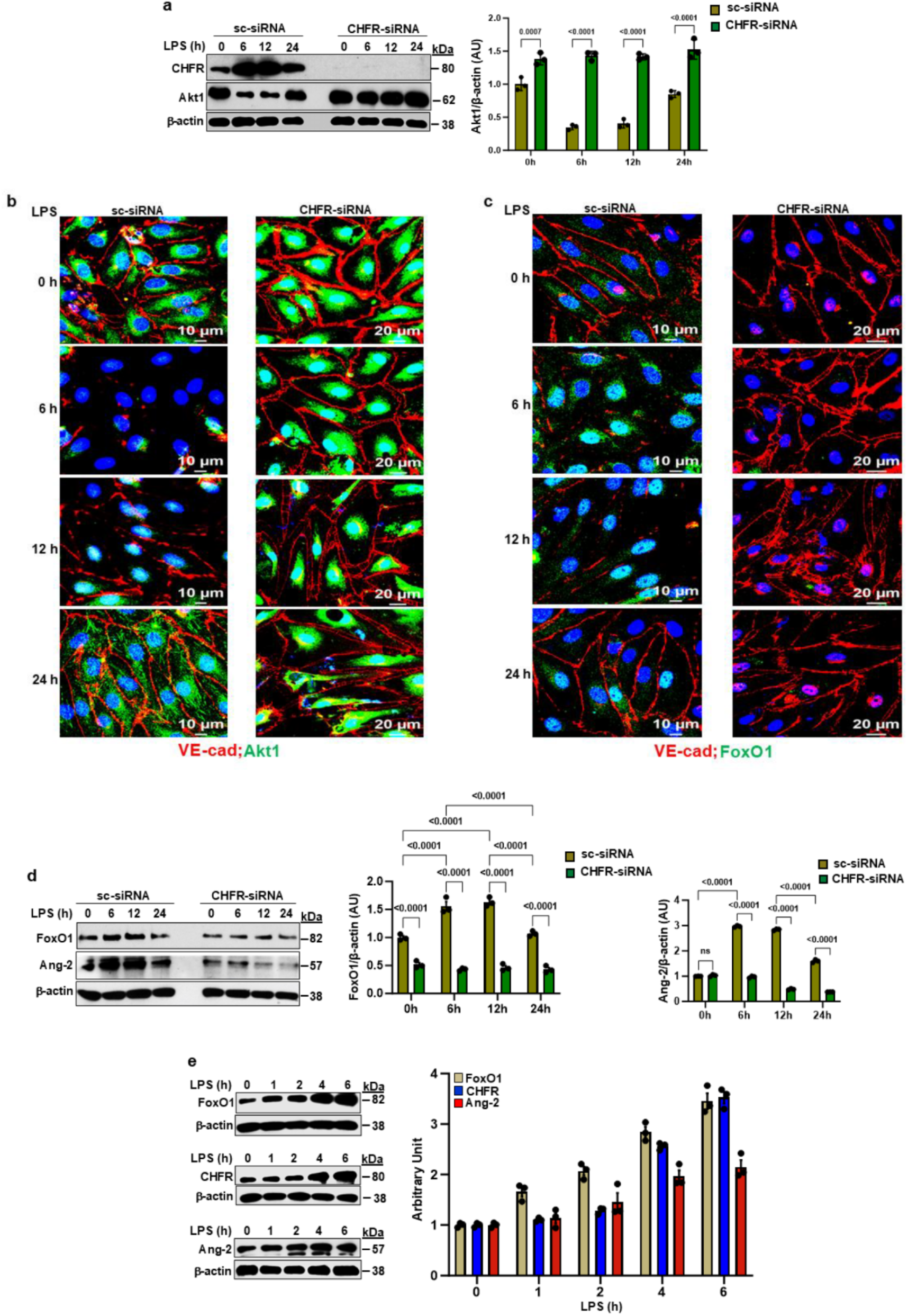
CHFR depletion in endothelial cells prevents LPS-induced downregulation of Akt1 and VE-cadherin, and the induction of FoxO1. **a** HLMVEC were transfected with Sc-siRNA or CHFR-siRNA. At 72 h post-transfection, cells challenged with LPS (5 μg/ml) for different time periods were used for IB analysis to determine expression of CHFR and Akt1. Shown are mean values ± SEM (*n* = 3 independent experiments; two-way ANOVA followed by Tukey’s post-hoc test). **b-c** HLMVEC were transfected with Sc-siRNA or CHFR-siRNA. At 72 h post transfection, cells challenged with LPS (5 μg/ml) for different time periods were stained with antibodies specific to VE-cadherin, Akt1, and FoXO1. Confocal imaging shows that CHFR depletion in EC prevents LPS-induced degradation of Akt1 and VE-cadherin and nuclear accumulation of FoxO1. **d** CHFR depletion prevents LPS-induced expression of FoxO1 and Ang-2. HLMVEC were transfected with Sc-siRNA or CHFR-siRNA as above and exposed to LPS for different time periods were used for IB to determine expression of FoxO1 and Ang-2. **e** LPS-induced time-course expression of FoxO1, CHFR, and Ang-2 in HLMVEC. HLMVEC exposed to LPS (5 μg/ml) for 0, 1, 2, 4, and 6 h were used for IB. **d-e** Shown are mean values ± SEM (*n* = 3 independent experiments; two-way ANOVA followed by Tukey’s post-hoc test).

Interestingly, LPS challenge increased nuclear presence of FoxO1 in control cells at 6 and 12 h, whereas FoxO1 expression and nuclear accumulation were blocked in CHFR-depleted cells (**Fig. 3c**). Immunoblotting confirmed increased FoxO1 expression in control HLMVEC following LPS exposure, an effect that was abolished by CHFR depletion (**Fig. 3d**). These data indicate that endothelial CHFR is required for LPS-induced expression of FoxO1 and Ang-2.

Because FoxO1 transcriptionally regulates CHFR expression in endothelial cells (*13*), we next examined whether CHFR stabilizes FoxO1 through a feed-forward mechanism. Time-course analysis revealed low basal expressions of FoxO1, CHFR, and Ang-2 in HLMVEC (**Fig. 3e**). FoxO1 expression was detectable within 1 h of LPS exposure and increased steadily at 2, 4, and 6 h. CHFR and Ang-2 expression were delayed and significantly elevated at 4 and 6 h (**Fig. 3e**). Consistently, pharmacological inhibition of Akt1 enhanced LPS-induced expression of FoxO1, Ang-2, and CHFR (**Supplemental Fig. 2a-c**) while reducing VE-cadherin expression (**Supplemental Fig. 2d**). These findings suggest that LPS-induced FoxO1 initially triggers CHFR expression, which subsequently downregulates Akt1, thereby stabilizing FoxO1 in a feed-forward loop.

### CHFR ubiquitylates phosphorylated Akt1

Since LPS exposure caused loss of Akt1 expression both *in vitro* and *in vivo*, we investigated whether CHFR regulates Akt1 stability through ubiquitylation. In control HLMVEC, LPS induced K^48^-linked polyubiquitylation of Akt1, whereas K^63^-linked ubiquitylation was not observed (**Fig. 4a**). Remarkably, K48-linked Akt1 ubiquitylation was abolished in CHFR-depleted cells following LPS challenge, while K^63^-linked ubiquitylation became detectable (**Fig. 4a**). Consistent with these findings, LPS induced K^48^-linked Akt1 ubiquitylation in lungs of *Chfr^fl/fl^* mice, whereas this modification was absent in *Chfr^ΔEC^* mice, despite the presence of K^63^-linked Akt1 ubiquitylation (**Fig. 4b**). These results indicate that LPS-induced Akt1 degradation occurs via CHFR-mediated K^48^-linked polyubiquitylation.

**Figure 4.**
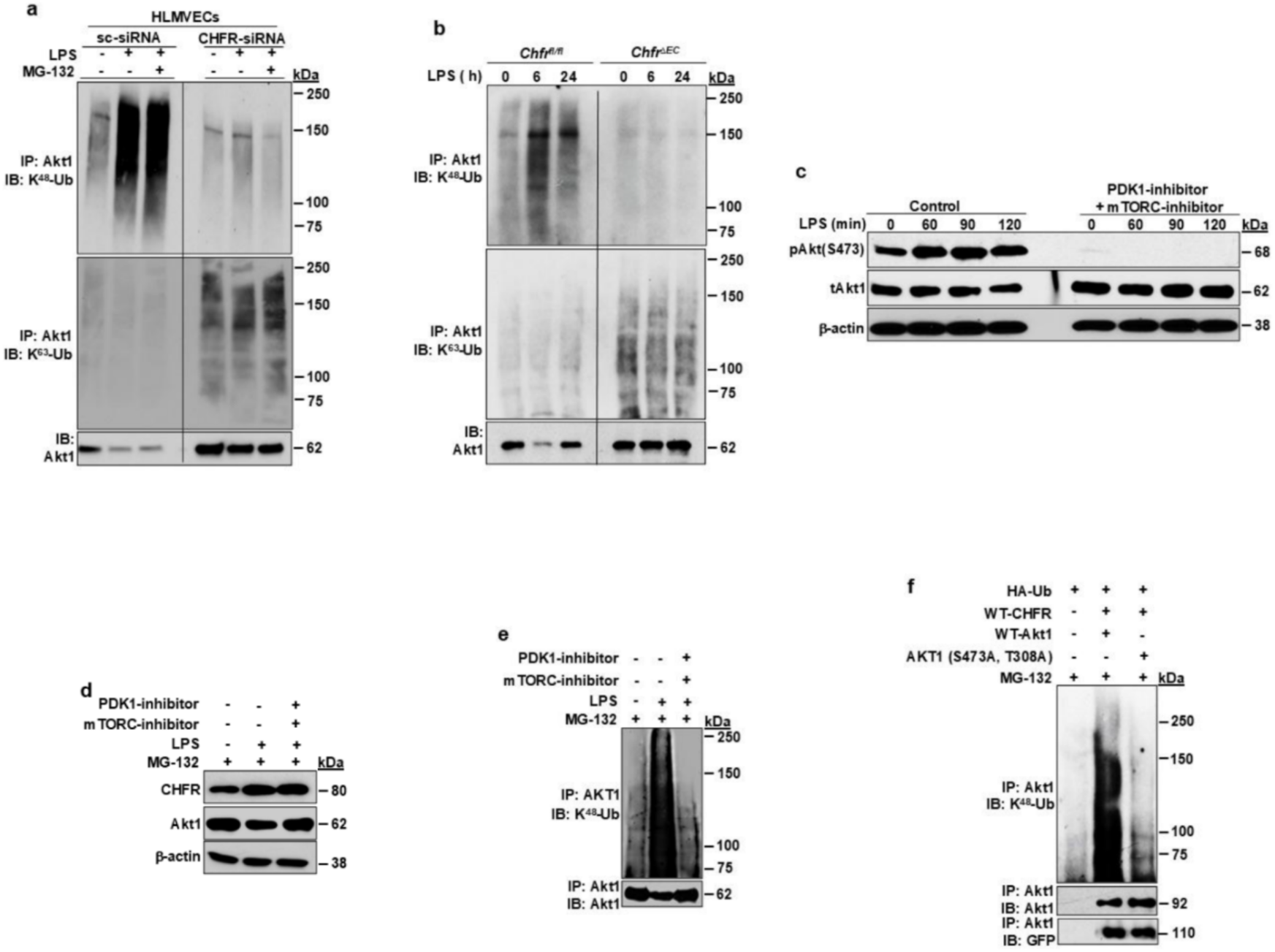
E3 ligase CHFR mediates ubiquitylation of phosphorylated Akt1. **a-b** CHFR deficiency in EC prevents LPS-induced ubiquitylation of Akt1. HLMVEC transfected with sc-siRNA or CHFR-siRNA. At 72 h post-transfection, cells were challenged with LPS (5 μg/ml) for 6 h in the presence of the proteasomal inhibitor MG132 (10 μM). Cell lysates were immunoprecipitated with anti-Akt1 mAb and blotted with antibodies specific to K^48^-linked poly-Ub or K^63^-linked poly-Ub (*n* = 2 independent experiments) (**a**). *Chfr^fl/fl^* (WT) and *Chfr^ΔEC^* mice were challenged with LPS (10 mg/kg; i.p.) for 6 h. After the LPS challenge, lungs harvested were used to determine ubiquitylation of Akt1 as above in **a (b)**. **c-f** CHFR induces ubiquitylation of activated Akt1. **c** Control and CHFR-depleted HLMVEC were pretreated with inhibitors of PDK1 and mTORC2 for 30 min and then exposed to LPS (5 μg/ml). Thereafter, cell lysates were used for IB to assess phosphorylation of Akt1. **d-e** HLMVEC pretreated with or without PDK1 and mTORC2 inhibitors for 1 h and stimulated with LPS (6 h) in the presence of MG123 (10 μM) showed that CHFR binds and ubiquitylates phosphorylated Akt1. **f** HEK-293T cells were transfected with HA-tagged ubiquitin (HA-Ub) (0.5 μg/ml) alone or co-transfected with N-terminal GFP-tagged WT-CHFR (1.5 μg/ml), N-terminal pmCherry-tagged WT-Akt1 (1.5 μg/ml), and phosphorylation-defective Akt1 mutant (Akt1T308A/S473A) (1.5 μg/ml) plasmids. Thirty-six hours after transfection, cells were incubated with MG123 (10 μM) for 3h, and cell lysates were used to determine phosphorylation-dependent ubiquitylation of Akt1.

CHFR is a RING domain-containing E3 ligase that preferentially targets phosphorylated substrates (*31*, *38*). Because Akt1 activation requires phosphorylation at T308 by PDK1 and S473 by mTORC2 (*39*), we examined whether Akt1 phosphorylation is required for CHFR-mediated ubiquitylation. Inhibition of PDK1/mTORC2 blocked both LPS-induced phosphorylation and ubiquitylation of Akt1 (**Fig. 4c-e**). Furthermore, LPS-induced K^48^-linked ubiquitylation of Akt1 was abolished when a phosphorylation-defective Akt1 mutant (Akt1-T308A/S473A) was expressed in TLR4-expressing HEK293 cells (**Fig. 4f**). These findings demonstrate that CHFR selectively targets phosphorylated AKT1 for degradation.

### CHFR binds and preferentially ubiquitylates Akt1 via K48-linked polyubiquitin chains

CHFR contains an N-terminal forkhead-associated (FHA) domain, a cysteine-rich (CR) domain, a RING finger (RF) domain, and a C-terminal poly-ADP-ribose–binding zinc finger (PBZ) motif (*13*, *38*). To determine whether CHFR directly interacts with Akt1, we co-expressed WT-Akt1 with WT or mutant CHFR constructs in TLR4-expressing HEK293 cells (**Fig. 5a-c**). Pull-down assays revealed that the RF domain of CHFR is essential for interaction with Akt1 (**Fig. 5c**).

**Figure 5.**
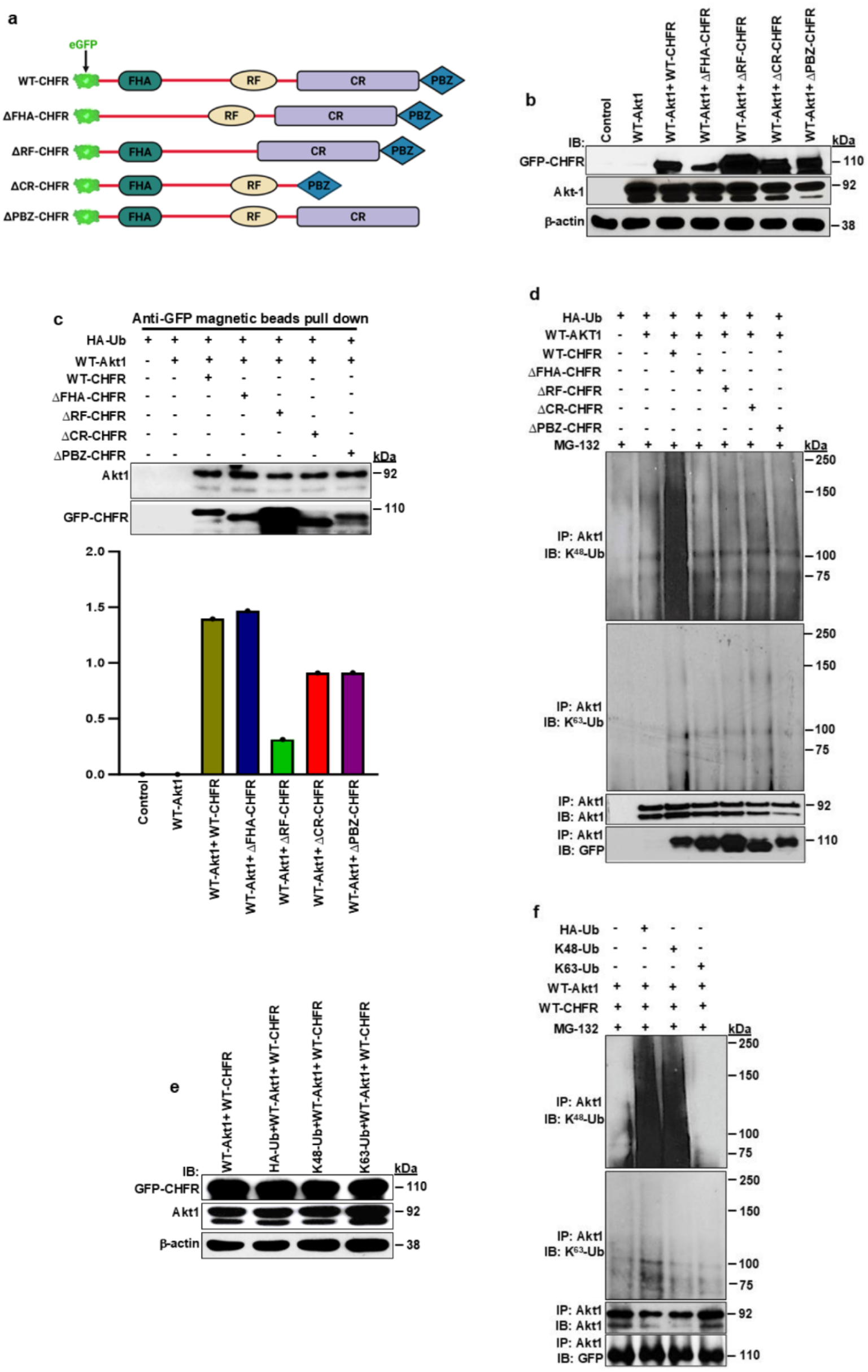
The ring finger domain of CHFR interacts with and mediates K^48^-linked polyubiquitylation of Akt1. **a** Schematics of the domain structures of human CHFR WT and mutants lacking forkhead-associated domain (ΔFHA-CHFR), RING finger domain (ΔRF-CHFR), cysteine-rich domain (ΔCR-CHFR), or poly-ADP ribose binding zinc-finger domain (ΔPBZ-CHFR) used in experiments. **b** Immunoblot showing expression of eGFP-tagged CHFR (WT) and CHFR mutants (1.5 μg/ml), along with pmCherry-tagged WT-Akt1 (1.5 μg/ml) in HEK-293T cells. **c** Transfected HEK-293T cells were used for anti-GFP agarose beads pull-down assays. Results show that WT-CHFR and CHFR mutants bind to WT-Akt1 *in vitro*. Bottom panel shows quantification of CHFR binding to Akt1 as a ratio of Akt1-to-GFP-CHFR. arb. units, arbitrary units. **d** HEK-293T cells transfected with WT-HA-Ub (0.5 μg/ml) alone or co-transfected with WT-CHFR (1.5 μg/ml), ΔFHA-CHFR (1.5 μg/ml), ΔRF-CHFR (1.5 μg/ml), and WT-Akt1 (1.5 μg/ml) were used to assess ubiquitylation of Akt1. At 36 h after transfection, cells were incubated with MG132 (10 μM) for 3 h, and then cell lysates were used for IB analysis. **e-f** HEK-293T cells transfected with WT-HA-Ub (0.5 μg/ml) or HA-tagged Ubiquitin where all Lysin (K) residues were mutated to Alanine (A) except at K48 or K63, along with WT-CHFR, and WT-Akt1 were used to determine CHFR-mediated linkage specific polyubiquitylation of Akt1.

Co-expression of WT-CHFR with WT-Akt1 resulted in robust K^48^-linked polyubiquitylation of AKT1, whereas deletion of any CHFR domain markedly reduced this modification (**Fig. 5d**). Notably, CHFR did not induce K^63^-linked Akt1 ubiquitylation (**Fig. 5d**). Further, linkage specificity was confirmed using ubiquitin mutants: CHFR-mediated Akt1 ubiquitylation occurred with WT or K^48^-only ubiquitin but not with K63-only ubiquitin (**Fig. 5e, f**). In other experiments, we co-expressed pmCherry-Akt1 (WT), eGFP-CHFR (WT), and HA-Ub (WT) in HEK293 cells and then confocal images were acquired after staining with K^48^-linkage specific polyubiquitin antibody. Here we observed that colocalization of K^48^-linked polyubiquitin in WT-CHFR and WT-AKT1 expressing cells (**Supplemental Figure 3**). These data together establish that CHFR promotes ubiquitylation of Akt1 through K^48^-linked polyubiquitin chains.

### Identification of Akt1 ubiquitylation sites targeted by CHFR

To identify Akt1 lysine residues targeted by CHFR, we performed mass spectrometry analysis (**Supplemental Fig. 4a**). HEK293 cells expressing mCherry-Akt1 and eGFP-CHFR were subjected to tryptic digestion, enrichment of ubiquitylated peptides using anti-di-glycine remnant antibodies (*40*), and then LC-MS/MS analysis (*41*). Four ubiquitylation sites “K30, K39, K154, and K268” were identified (**Supplemental Fig. 4a-c**). To validate these sites, we generated Akt1 lysine-to-arginine (K/R) mutants and co-expressed them with HA-ubiquitin and WT-CHFR. CHFR-induced K^48^-linked polyubiquitylation was markedly reduced in Akt1-K39R and Akt1-K268R mutants and completely abolished in double (K39R/K268R) and quadruple (K30R/K39R/K154R/K268R) mutants (**Supplemental Fig. 4d**). These findings indicate that CHFR targets lysine residues in both the N-terminal and kinase domains of Akt1 (**Supplemental Fig. 4c)** to induce proteasomal degradation.

### Ubiquitylation-defective Akt1 mutants prevent LPS-induced endothelial barrier breakdown in vitro *and* in vivo

To assess the functional significance of CHFR-mediated Akt1 degradation, we expressed WT or K/R mutant Akt1 constructs in human dermal microvascular endothelial cell line (HMEC). LPS induced downregulation of Akt1 and VE-cadherin in WT-Akt1-expressing cells, whereas this effect was prevented in cells expressing Akt1-K39R/K268R or Akt1-K30R/K39R/K154R/K268R (**Fig. 6a**). Immunofluorescence analysis confirmed preservation of VE-cadherin at AJs in cells expressing ubiquitylation-defective Akt1 mutants (**Fig. 6b**). To evaluate *in vivo* relevance, WT-Akt1 or mutant Akt1 constructs were delivered to lung endothelial cells of WT mice via liposome-mediated transduction (*13*, *42*, *43*). We noted the expression of WT-Akt1 or mutant Akt1 in intact lung microvessels (**Fig. 6c**). LPS challenge induced degradation of VE-cadherin in control and WT-Akt1-expressing mice, whereas this effect was abolished in mice expressing ubiquitylation-defective Akt1 mutants (**Fig. 6d**). Consistently, LPS-induced lung vascular leak (**Fig.6e**) and neutrophil sequestration (**Fig. 6f**) were significantly reduced in mutant Akt1-expressing mice compared with controls (**Fig. 6e, f**). Together, these findings demonstrate that CHFR-mediated K^48^-linked polyubiquitylation and degradation of Akt1 is a critical mechanism driving endothelial barrier breakdown during inflammatory stress. Based on the present study results, we propose model for the regulation of endothelial junctional barrier under basal and inflammatory conditions such as sepsis (**Fig. 6g**).

**Figure 6.**
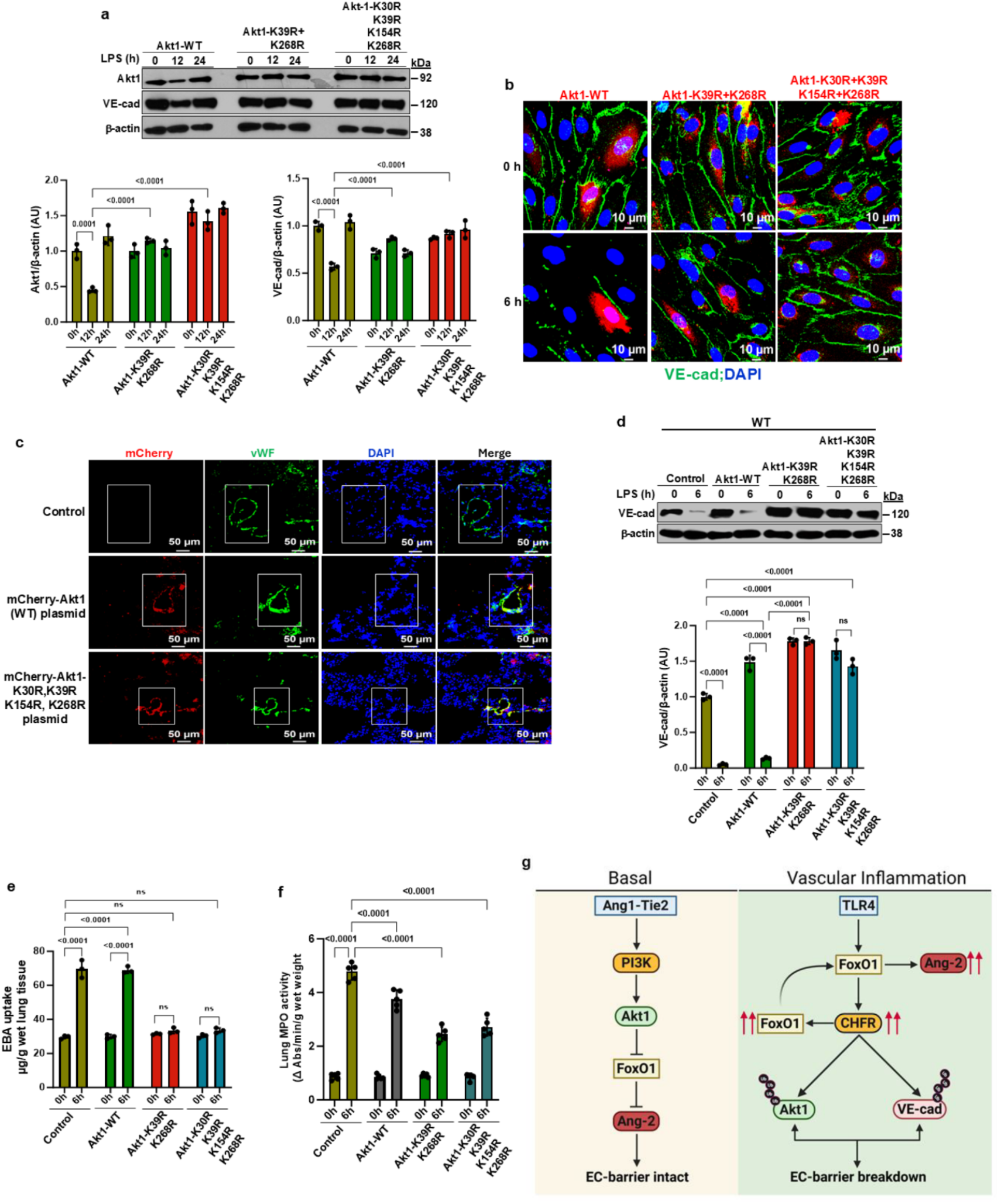
Expression of a ubiquitylation-defective Akt1 mutant prevents LPS-induced degradation of Akt1 and VE-cadherin in EC and mitigates lung vascular leak *in vitro* and *in vivo*. **a** HMEC were transfected with WT or K/R-mutant Akt1 constructs. At 36 h after transfection, cells were treated with LPS (5 μg/ml) for 0, 12, and 24h and then cells were used for IB to determine expression of Akt1 and VE-cadherin. Shown are mean values ± SEM (n = 3 independent experiments; two-way ANOVA followed by Tukey’s post hoc test). **b** TIME endothelial cells (telomerase-immortalized human dermal microvascular endothelial cell line) were transfected with WT or K/R mutant Akt1 constructs and stimulated with LPS (5 μg/ml) for 0 and 6 h. Confocal imaging showed that expression of K/R mutant Akt1 prevents degradation of VE-cadherin. **c** WT mice were injected (i.v.) with liposome-encapsulated pmCherry-tagged WT or K/R mutant Akt1 constructs. Lungs harvested 96 h after injection were subjected to cryosection and stained with EC marker antibody vWF (green). Confocal imaging confirms expression of pmCherry-Akt1 (red) plasmid in lung endothelial cells. **d-f** Liposome-mediated delivery of Akt1 (WT) or K/R-mutated Akt1 in WT mice prevents degradation of VE-cadherin, mitigates LPS-induced lung vascular leak (EBA uptake), and reduces PMN transmigration (MPO assay). Shown are mean values ± SEM (*n* = 3 or *n* = 5 mice/group; two-way ANOVA followed by Tukey’s post hoc test). **g** Model for E3 ligase CHFR regulation of endothelial junctional barrier integrity. Under baseline condition, constitutive Ang1-Tie2 signaling in EC maintains endothelial junctional barrier through Akt1 activation-mediated inhibition of the transcription factor FoxO1 activation and Ang-2 expression. During vascular inflammatory conditions such as sepsis, TLR4 signaling induces the expression of E3 ligase CHFR in a FoxO1-dependent manner. Then the upregulated CHFR mediates K^48^-linked polyubiquitylation and degradation of Akt1 and VE-cadherin (Tiruppathi et al., 2023) to disassemble EC junctional barrier. CHFR-mediated loss of FoxO1 negative regulator Akt1 expression in EC leads to increased FoxO1 expression which in turn promotes sustained expression of Ang-2 in EC to induce life-threatening pulmonary edema.

## Discussion

The Ang-1/Tie2-Akt1 axis is constitutively active under basal conditions and preserves endothelial junctional barrier integrity (*7*, *25*). Under systemic inflammatory conditions such as sepsis, Ang-1/Tie2-Akt1 axis signaling becomes dysregulated, whereas FoxO1-mediated expression of the Tie2 antagonist Ang-2 is elevated, leading to compromised endothelial barrier function (*7*). Although FoxO1-induced Ang-2 expression promotes endothelial barrier breakdown, resulting in pulmonary edema and vascular remodeling (*44*), little is known about the intrinsic signaling pathway that modulates endothelial Ang-2 production during vascular inflammation. In the present study, we demonstrated that TLR4 signaling–induced expression of the E3 ligase CHFR represents a key mechanism driving K^48^-linked polyubiquitylation and proteasomal degradation of Akt1, thereby impairing Tie2 signaling in endothelial cells. Gene knockdown and EC-restricted knockout mouse model studies have established the role of CHFR in regulating Akt1 function and, consequently, endothelial junctional barrier integrity. Loss of CHFR in endothelial cells augmented Akt1 expression and activity and suppressed FoxO1 and its target gene Ang-2 expression. Together, the findings of the present study uncover a critical role for CHFR in regulating Tie2 signaling through ubiquitylation-mediated degradation of Akt1 in endothelial cells.

Because EC-expressed Akt1 plays a crucial role in regulating endothelial junctional barrier stability (*3*, *33*, *45*), we investigated whether the E3 ligase CHFR modulates Akt1 expression and function in EC. Notably, we observed increased Akt1 expression accompanied by decreased expression of FoxO1 and its target gene Ang-2 in EC-restricted *Chfr* knockout (*Chfr^ΔEC^*) mice. Importantly, LPS-induced Akt1 degradation was prevented in *Chfr^ΔEC^*mice. In support of these observations, LPS-induced Ang-2 and FoxO1 expression was also blocked in *Chfr^ΔEC^*mice. Furthermore, we found increased expression of Ang-1 and Tie2 in *Chfr^ΔEC^* mice. Consistent with the mouse studies, depletion of CHFR in human lung microvascular endothelial cells (HLMVEC) increased Akt1 expression, and in these CHFR-depleted cells, LPS-induced downregulation of Akt1 was prevented. Remarkably, Ang-1 stimulation induced robust and sustained phosphorylation of Akt1 at S473 and T308 in CHFR-deficient EC compared with control EC. Using Akt FRET biosensors in live endothelial cells, we also demonstrated that CHFR depletion increased both basal and Ang-1-induced Akt1 activity. In addition, Ang-1-induced Tie2 phosphorylation and phosphorylation of GSK3β at S9 were sustained in CHFR-deficient HLMVEC. Collectively, these results support the proposal that CHFR-mediated downregulation of Akt1 may contribute to endothelial barrier dysfunction through enhancing FoxO1 function.

Although Akt1 expression and function can be regulated by post-translational modification through ubiquitylation (*28*), little is known about whether Akt1 expression and function are directly regulated by ubiquitylation in EC. Therefore, we examined whether the E3 ligase CHFR promotes Akt1 degradation via ubiquitylation in response to LPS. We observed that LPS-induced ubiquitylation of Akt1 through K^48^-linked polyubiquitin chains was blocked in CHFR-deficient human lung EC and in EC-restricted *Chfr* knockout (*Chfr^ΔEC^*) mice. Furthermore, we demonstrated that CHFR interacts with and ubiquitylates phosphorylated Akt1. Using phosphorylation-defective Akt1 mutants, we showed that CHFR failed to induce ubiquitylation of phosphodeficient Akt1. These findings are consistent with previous studies demonstrating that phosphorylated Akt1 is a preferred substrate for E3 ligases in tumor cell lines and fibroblasts (*29–31*). Thus, our results suggest that the E3 ligase CHFR regulates EC function through ubiquitylation of Akt1.

To determine the specificity of CHFR mediating ubiquitylation of Akt1, we performed *in vitro* ubiquitylation assays and observed that CHFR binds to and ubiquitylates Akt1. We also demonstrated that the RING domain of CHFR is required for Akt1 ubiquitylation. Additional experiments confirmed that CHFR-mediated ubiquitylation occurs through K^48^-linked, but not K^63^-linked, polyubiquitin chains. This finding is consistent with the concept that CHFR-mediated ubiquitylation of Akt1 via K^48^-linked polyubiquitin chains promotes proteasomal degradation. Using an LC-MS approach, we identified the critical lysine residues targeted by CHFR. Specifically, CHFR ubiquitylates Akt1 at “K30, K39, K154, and K268” residues. To assess the specificity of these lysine residues, we generated Akt1 K-to-R mutants and co-expressed them with HA-ubiquitin and WT-CHFR. CHFR-induced K48-linked polyubiquitylation was reduced in Akt1-K39R and Akt1-K268R mutants and completely abolished in double (K39R/K268R) and quadruple (K30R/K39R/K154R/K268R) mutants. These results indicate that CHFR targets lysine residues in both the N-terminal and kinase domains of Akt1 to promote its proteasomal degradation.

To determine the functional relevance of CHFR-mediated ubiquitylation of Akt1 in regulating endothelial junctional barrier integrity, we expressed WT-Akt1 or K/R mutant Akt1 in EC both *in vitro* and *in vivo*. Expression of WT or K/R mutant Akt1 constructs in endothelial cells showed that LPS induced downregulation of Akt1 and VE-cadherin, whereas this effect was blocked in cells expressing Akt1-K39R/K268R or Akt1-K30R/K39R/K154R/K268R. To assess *in vivo* relevance, WT-Akt1 or mutant Akt1 constructs were delivered to lung endothelial cells of WT mice via liposome-mediated transduction, and the effects of LPS on endothelial barrier stability were examined. LPS challenge induced degradation of VE-cadherin in control and WT-Akt1–expressing mice, whereas this effect was markedly reduced in mice expressing ubiquitylation-defective Akt1 mutants. In addition, LPS-induced lung vascular leak and neutrophil sequestration were significantly reduced in mutant Akt1–expressing mice compared with controls. Taken together, these findings support that CHFR-mediated K^48^-linked polyubiquitylation and degradation of Akt1 represent a critical mechanism driving endothelial barrier breakdown during inflammatory stress.

The angiopoietin-1/2–Tie2 axis is crucial for systemic regulation of endothelial activation and dysfunction in various pathological conditions associated with systemic inflammation (*7*, *46*). Ang-1 stabilizes the endothelium, whereas Ang-2 destabilizes it; however, the regulation of Ang-1 and Ang-2 expression has remained poorly understood. In this study, we showed that CHFR deficiency increased Ang-1 levels, which may be attributable to enhanced stability of EC–pericyte interactions. Importantly, LPS-induced expression of Ang-2, a FoxO1 target gene, was blocked in *Chfr^ΔEC^*mice, whereas Akt1 expression was increased in these mice. Akt1 functions as a negative regulator of FoxO1 by promoting its phosphorylation and degradation. Thus, increased Akt1 expression could suppress FoxO1 expression and subsequent induction of its target gene Ang-2. The results of the present study support the concept that, during vascular inflammation, FoxO1-induced CHFR expression promotes degradation of Akt1, which in turn stabilizes FoxO1 expression and activity, leading to induction of the endothelial barrier–destabilizing Tie2 antagonist Ang-2.

In summary, we identified that vascular inflammation induced CHFR expression as a key mechanism that promotes K^48^-linked polyubiquitylation and degradation of Akt1 in endothelial cells. Accordingly, CHFR deficiency increased Akt1 and VE-cadherin expression in endothelial cells. Mechanistically, CHFR interacts with phosphorylated Akt1 and mediates its ubiquitylation at lysine residues K30, K39, K154, and K268. Expression of ubiquitylation-deficient Akt1 mutant prevented LPS-induced VE-cadherin degradation and vascular injury. Furthermore, in CHFR deficient endothelial cells, LPS-induced expression of FoxO1 and Ang-2 was prevented. Collectively, the present study identifies the E3 ligase CHFR as a critical regulator of endothelial inflammatory responses by controlling Akt1 stability and VE-cadherin expression during inflammation.

## Methods

### Primary Antibodies

Rabbit monoclonal antibody (mAb) against Akt1 (catalog #2938S; immunoblot [IB], 1:1000; immunostaining [IS], 1:150, immunoprecipitation [IP], 1 μg/100 μg cell lysate protein), rabbit mAb against FoxO1 (catalog #2880; IB, 1:1000; IS, 1:100), rabbit (mAb) against Phospho-FoxO1(T24) (catalog #9464; IB, 1:1000), rabbit (mAb) against Phospho-FoxO1(S256) (catalog #9461; IB, 1:1000), rabbit (mAb) against GSK3β (catalog #9315; IB, 1:1000), rabbit (mAb) against Phospho-GSK3β (catalog #9322; IB, 1:1000), rabbit mAb against Akt1 (catalog #2938; IB, 1:1000; IS, 1:100), rabbit (mAb) against Phospho-Akt1(T308) (catalog #13038; IB, 1:1000), rabbit (mAb) against Phospho-Akt1(S473) (catalog #4058; IB, 1:1000), rabbit (mAb) against phospho-β-catenin (catalog #9561; IB, 1:1000), rabbit mAb against ubiquitin K63-linkage-specific (catalog #5621S; IB, 1:1000), rabbit mAb against ubiquitin K48-linkage-specific (catalog #4289S; IB, 1:1000 were from Cell Signaling Technology. Mouse mAb against Akt1 (catalog #sc-5298; IB, 1:1000; IP, 1 μg/100 μg cell lysate protein), and mouse mAb against FoxO1 (catalog #sc-374427; IS, 1:100) were from Santa Cruz Biotechnology. Mouse mAb against Tie2 (catalog #05-584; IB, 1:1000) was from Millipore Sigma. Rabbit polyclonal antibody (pAb) against phospho-Tie2 (catalog #AF3909; IB, 1:1000), goat polyclonal antibody (pAb) against PDGFRβ (catalog #AF1042; IS, 1:100) were from R&D system. Mouse mAb against β-actin (catalog #A5441; IB, 1:2000) was from Sigma-Aldrich Inc. Rabbit pAb against GFP (catalog #50430-2-AP; IB, 1:2000), rabbit pAb against CHFR (catalog #12169-1-AP; IB, 1:1000), rabbit pAb against Angiopoietin-2 (catalog #24613-1-AP; IB, 1:1000), rabbit pAb against β-catenin (catalog #61067-2-AP; IB, 1:1000), and rabbit pAb against mCherry (catalog #26765-1-AP; IB, 1:1000) were from Proteintech. Rabbit polyclonal antibody (pAb) against the C-terminus of VE-cadherin (catalog #Ab33168; IB, 1:1000, IS, 1:150), rat mAb against CD31 (catalog #Ab7388; IS, 1:150), and sheep pAb against vWF (catalog #ab11723; IS, 1:150) were from Abcam.

### Secondary antibodies

Anti-rabbit IgG (H + L) peroxidase labeled (catalog #5220-0458; IB, 1:5000) and anti-mouse IgG (H + L) peroxidase labeled (catalog #5450-0011; IB, 1:5000) were from Sera Care KPL. Donkey anti-goat IgG (H + L) HRP labeled (catalog #A16005; IB, 1:5000), chicken anti-rabbit IgG (H + L) Alexa Fluor 594 labeled (catalog # A21442; IS, 1:500), chicken anti-mouse IgG (H + L) Alexa Fluor 647 labeled (catalog #A21463; IS, 1:500), chicken anti-mouse IgG (H + L) Alexa Fluor 488 labeled (catalog #A21200; IS, 1:500), goat anti-rabbit (H + L) Alexa Fluor 488 labeled (catalog #A11034; IS, 1:500), donkey anti-sheep IgG (H + L) Alexa Fluor 488 labeled (catalog #A11015; IS, 1:500), rabbit anti-goat IgG (H + L) Alexa Fluor 488 labeled (catalog # A11078; IS, 1:500), rabbit anti-rat IgG (H + L) Alexa Fluor 594 labeled (catalog # A21211; IS, 1:500), chicken anti-rabbit IgG (H + L) Alexa Fluor 488 labeled (catalog #A21441; IS, 1:500), and chicken anti-rabbit IgG (H + L) Alexa Fluor 594 labeled (catalog #A21442; IS, 1:500) were from Invitrogen.

### Chemicals, siRNAs, peptides, and expression plasmids

Lipopolysaccharide (E. coli O111:B4; catalog # L3012) was from Sigma-Aldrich Inc. Recombinant human angiopoietin-1 (catalog #923-AN-025) was from biotechne (R&D system). Akt1/2 inhibitor (catalog #A6730) was from Millipore Sigma. Mammalian targets of rapamycin (mTOR) inhibitor (catalog #WYE-354), and 3-Phosphoinositide-Dependent Kinase 1 (PDK1) inhibitor (catalog #GSK2334470) were from Selleckchem.com.

Scrambled-siRNA (Sc-siRNA) and human (*h*)-specific siRNA1 against CHFR (A pool of 3 target-specific siRNAs, sense: 5′-CUCUGUGGCAAGUGAUGAAtt-3′; sense: 5′-GAAGAAGAUGUGCAAAGUAtt; sense: 5′-GGUAUAGUAUGGUAUUUGAtt-3′) were obtained from Santa Cruz Biotechnology. pEGFP-C2 plasmid (Addgene catalog #61853) encoding wild-type human CHFR (WT*h*-CHFR), *h*CHFR mutant lacking forkhead-associated domain (ΔFHA-CHFR), *h*CHFR mutant lacking RING finger domain (ΔRF-CHFR), mutant lacking cysteine-rich domain (ΔCR-CHFR), and mutant lacking poly-ADP ribose binding zinc-finger domain (ΔPBZ-CHFR) were custom prepared by Genscript (Piscataway, NJ). pmcherry-C1 plasmid (Addgene catalog #86631) encoding wild-type human Akt1 (WT *h*-Akt1), phopsho-defective *h*Akt1 (T308A and S473A) mutant and ubiquitylation-defective *h*Akt1 (K30R, K39R, K39R+154R and K30R+K39R+K154R+K268R) mutants were custom prepared by Genscript (Piscataway, NJ). HA-tagged WT-ubiquitin-expressing plasmid (catalog #17608) and HA-tagged K48- or K63-ubiquitin-expressing plasmids (where all the lysine residues were mutated to Arginine except K48 or K63) were from Addgene. CHFR deletion constructs and Akt1 mutant constructs were prepared using standard molecular biology methods. DNA sequencing was performed to verify the deletion. PCR primers were custom synthesized by Integrated DNA Technologies (Coralville, IA).

### Cells

Human lung microvascular endothelial cells (HLMVEC; Lonza, catalog#CC-2527) were utilized between passages 3 and 7 for experiments. The telomerase-immortalized human dermal microvascular endothelial cell line (Lonza, catalog #CRL-4025) was maintained in Vascular Cell Basal Medium (ATCC^®^ PCS-100-030), supplemented with Microvascular Endothelial Cell Growth Kit-VEGF (ATCC^®^ PCS-110-041). The human dermal microvascular endothelial cell line (HMEC), kindly provided by Dr. Thomas J. Lawley (Emory University), was maintained in MCDB131 medium supplemented with 10% FBS, 10 ng/ml epidermal growth factor, 2 mM L-glutamine, and 1 μg/ml hydrocortisone. For gene silencing experiments, HLMVEC and human dermal microvascular endothelial cell line were grown to approximately ∼80% confluence on gelatin-coated plates and transfected with either target-specific or scrambled control (sc-siRNA). Experimental procedures were performed after 72 hours of post-transfection.

### Mice

All Animal procedures were conducted in accordance with protocols approved by the Office of Animal Care and Institutional Biosafety (OACIB) at the University of Illinois at Chicago. Mice were housed in a pathogen-free environment under a 12 h light/dark cycle at 23 °C and 40–60% humidity, with *ad libitum* access to food and water. Both male and female mice (8–12 weeks old) were utilized, with specific cohort sizes detailed in the figure legends. *Chfr^flox/flox^* mice (C57BL/6 background) were generated using a targeting vector to flank exon 3 of the *Chfr* gene with loxP sites as we described previously (Tiruppathi et al., 2023). To generate endothelial cell (EC)-restricted *Chfr* knockout (*Chfr^ΔEC^*) mice, *Chfr^flox/flox^* (*Chfr^fl/fl^*) mice were crossed with B6.Cg-Tg (Cdh5-cre)7Mlia/J (VE-cadh-Cre^+^) transgenic mice from Jackson Laboratory.

### Akt activity measurement using FRET Biosensors

The Akt FRET biosensor utilized to assess live cell AKT1 activity was previously reported (*35*). The biosensor contains both an Akt substrate and a phospho-peptide binding domain. When the substrate becomes phosphorylated by endogenous Akt kinase and the domain subsequently binds the phosphorylated substrate, the efficiency of Forster Resonance Energy Transfer (FRET) between the two fluorescent proteins within the biosensor construct increases. To detect the increase in FRET, fluorescence microscopy was performed using excitation bands 510/25 and 550/15 nm and emission bands 540/21 and 595/40 nm for YFP and RFP, respectively. Intensity images were taken for tens of fields-of-view over 60 minutes at 30 seconds interval. The background-subtracted FRET ratio in each single cell was used to quantify the kinetic Akt activity level. Control HMEC and CHFR-siRNA transfected HMEC were used for experiments to assess basal as well as rhAng-1 stimulated Akt activity. Forty-eight hours after transfection of Sc-siRNA or CHFR-siRNA, cells were transfected Akt FRET biosensor plasmid and at 24 hours, cells were used to measure live cell activity.

### Plasmid transfections and anti-GFP antibody bead pull-down assay

HEK293 cells (ATCC, catalog #CRL-1573) grown to ∼80% confluency were transfected with HA-tagged ubiquitin (HA-Ub) (0.5 μg/ml) alone or co-transfected with plasmids encoding N-terminal eGFP fused WT-*h*CHFR (1.5 μg/ml), or ΔFHA-*h*CHFR (1.5 μg/ml), or ΔRF-*h*CHFR (1.5 μg/ml) ΔCR-*h*CHFR (1.5 μg/ml), or ΔPBZ-*h*CHFR (1.5 μg/ml) as well as plasmids encoding N-terminal mCherry fused WT-*h*Akt1 (1.5 μg/ml) using CalPhos mammalian transfection kit from ClonTech laboratories. WT-*h*CHFR or mutant CHFRs binding to WT-*h*Akt1 was determined by anti-GFP antibody agarose beads (GFP-Trap Agarose, Chromotek catalog #gta-10) pull down as we described previously (*13*). At 48 hours of post-transfection, cells were washed twice with ice-cold PBS and then lysed with RIPA lysis buffer (50 mM Tris-HCl, pH7.5, 150 mM NaCl, 1 mM EGTA, 1% Triton X-100, 0.25% sodium deoxycholate, 0.1% SDS, 10 μM orthovanadate, and protease-inhibitor mixture) for 30 min at 4^0^C. Cell lysates were centrifuged (30,000 × *g* for 10 min) to remove insoluble materials. Clear supernatant collected containing 150 μg protein was incubated with 30 μl anti-GFP agarose beads (ChromTech, catalog #gta-10) for 2 h at 4 °C on a rotating platform. Thereafter, the beads were briefly washed 4 times with wash buffer (10 mM Tris-HCl, pH7.5, 150 mM NaCl, 0.1% Triton X-100) by centrifugation at 500 × *g* for 5 min and then used for IB analysis.

### Ubiquitylation assay

HEK293 cells (ATCC, catalog#CRL-1573) were transfected according to the previously described protocol (*13*). At 48 hours of post-transfection, cells were treated with the proteasome inhibitor MG132 (10 μM) for 4 hours to stabilize ubiquitinated proteins. Cell lysates were then subjected to immunoprecipitation using a monoclonal antibody (mAb) against Akt1. The resulting precipitates were analyzed via immunoblotting with linkage-specific antibodies directed against K48-linked or K63-linked ubiquitin chains.

To detect ubiquitination of Akt1 by the E3 ligase CHFR by confocal imaging, we transfected the HEK-293T cells with HA-tagged WT ubiquitin (HA-Ub) (0.5 μg/ml) alone or co-transfected with GFP-tagged WT-CHFR (1.5 μg/ml), and pmCherry-tagged WT-Akt1 (1.5 μg/ml). After 24 h of transfection, the cells were trypsinized and seeded on a glass cover slip. After 24 h, the cells were fixed in 4% paraformaldehyde (PFA) for 15 minutes at 4 °C. Following a 1-minute permeabilization step with 0.05% Triton X-100 in PBS, cells were blocked for 1 hour in a solution of PBS containing 5% horse serum and 1% BSA. Cells were incubated with an anti-HA primary antibody overnight at 4 °C in PBS containing 1% BSA. After subsequent washes, samples were incubated with Alexa-Fluor conjugated secondary antibodies for 1 hour at room temperature. Coverslips were then mounted using an anti-fade medium containing DAPI (Invitrogen). The images were acquired using Zeiss LSM 880 confocal microscopy.

### Liposome-mediated plasmid delivery

Liposomes were prepared as described previously (*13*, *42*, *43*). Briefly, a 1:1 molar mixture of dimethyldioctadecylammonium bromide (DDAB) and cholesterol was evaporated to dryness using a Rotavaporator (Brinkmann) and reconstituted in 5% dextrose. The solution was sonicated for 20 minutes and passed through a 45 μm syringe filter. mCherry-tagged WT-Akt1 or ubiquitination-defective Akt1 plasmids were then encapsulated at a ratio of 1 μg DNA to 8 nmol liposomes. The liposomes and plasmid mixture were injected i.v. into WT mice (30 μg of plasmid in 150 μl suspension/mouse). Mice were used for experiments 96 hours post-delivery of liposomes/plasmid mixture.

### Lung injury measurements

The lung vascular leak was assessed according to the protocol described (*13*). Briefly, mice were injected (i.v., retro-orbital) with Evans blue dye-bound albumin (EBA, 1%) 30 min before sacrifice and lung perfusion for EBA measurement. Neutrophil infiltration was measured by myeloperoxidase (MPO) activity. Briefly, lung tissues were homogenized in 1 mL of 50 mM PBS (pH 6.0) containing hexadecyltrimethylammonium bromide (5 mg/mL), followed by sonication. The lysates were centrifuged at 20,000 ×g for 20 min, and the supernatant was collected. The supernatant was mixed with reaction buffer (0.2 mg/mL o-dianisidine hydrochloride and 0.0005% H2O2), and MPO activity was measured as a change in absorbance at 460 nm over 3 min.

### Confocal Microscopy

HLMVEC or HMEC grown to confluency on glass coverslips challenged with or without LPS (5 μg/ml) were briefly rinsed with ice-cold PBS and fixed in 4% paraformaldehyde (PFA) for 15 minutes at 4 °C. Following a 1-minute permeabilization step using 0.05% Triton X-100 in PBS, cells were blocked for 1 hour in a solution of PBS containing 5% horse serum and 1% BSA. Specific primary antibodies were applied overnight at 4 °C in PBS containing 1% BSA. After subsequent washes, samples were incubated with Alexa-Fluor conjugated secondary antibodies for 1 hour at room temperature. Coverslips were then mounted using an anti-fade medium containing DAPI (Invitrogen).

### Immunostaining of lung tissue sections

Formalin fixed, paraffin-embedded sections were deparaffinized using a graded xylene and ethanol series, then rehydrated in distilled water. Heat-induced epitope retrieval was performed in a citrate buffer (10 mM citrate, 0.05% Tween 20, pH 6.0) for 20 minutes. Following a 2-hour blocking step in a blocking buffer (PBS containing 4% BSA, 0.2% Triton X-100) at room temperature, sections were incubated overnight with primary antibodies. Immunostainings were detected using fluorochrome-conjugated secondary antibodies for 1 hour at room temperature. Finally, slides were mounted using an anti-fade medium containing DAPI and imaged via Zeiss LSM 880 confocal microscope.

### Immunoprecipitation and immunoblotting

HLMVEC and HMEC grown to confluence, treated with or without specific agents, were three times washed with phosphate-buffered saline (PBS) at 4 °C and lysed in lysis buffer. Mouse lungs were homogenized in lysis buffer. Endothelial cell lysates or mouse lung homogenates were centrifuged (30,000 × *g* for 10 min) to remove insoluble materials. Clear supernatant collected 50 μg protein, was subjected to immunoblot analysis. For immunoprecipitation, clear supernatant 150-300 μg of protein was used. Each sample was incubated overnight with indicated antibody at 4 °C on rotation. Next day, Protein A/G beads were added to the sample and incubated at 4 °C for 1 h on rotation. Protein A/G beads were then washed three times with wash buffer (Tris-buffered saline containing 0.05% Triton X-100, 1 mM Na_3_VO_4_, 1 mM NaF, 2 μg/ml leupeptin, 2 μg/ml pepstatin A, 2 μg/ml aprotinin, and 44 μg/ml phenylmethylsulfonyl fluoride) and used for immunoblotting analysis. Endothelial cell lysates, lung tissue extracts, or immunoprecipitated proteins were boiled in Laemmli sample buffer (50 mM Tris-HCl pH 6.8, 2% SDS, 10% glycerol, 0.05% bromophenol blue, and 2.5% β-mercaptoethanol) and resolved by SDS-PAGE on a 4-15% gradient or 10% separating gel and transferred to a polyvinylidene difluoride (PVDF) membrane. For the detection of ubiquitylation of proteins, immunoprecipitated proteins were boiled in sample buffer (100 mM Tris-HCl pH 6.8, 4% SDS, 20% glycerol, 0.2% bromophenol blue, and 200 mM DTT) and then proteins were resolved by SDS-PAGE on a 6% separating gel. Membranes were blocked with 5% dry milk in TBST (10 mM Tris-HCl pH7.5, 150 nM NaCl, and 0.05% Tween-20) at RT for 1 h. Membranes were then probed with the indicated primary antibody (diluted in blocking buffer) overnight at 4 °C. Next, membranes were washed 3 times and then incubated with appropriate HRP-conjugated secondary antibody. Protein bands were detected by enhanced chemiluminescence. Immunoblot bands were quantified using NIH ImageJ software.

### Mass spectrometry

The post-translational modification (ubiquitylation of Akt1) was identified as per the protocol provided by PTMScan® HS Ubiquitin/SUMO Remnant Motif (K-ε-GG) Kit (#59322, Cell Signaling Technology). Briefly, HEK-293T cells were transfected with pmCherry-Akt1 (WT), eGFP-CHFR (WT), and HA-Ub (WT). At 36h of transfection, cells were collected and lysed with urea lysis buffer (9 M urea, 20 mM HEPES, pH 8.0, 2X phosphatase inhibitor cocktail; 1 mM sodium orthovanadate, 2.5 mM sodium pyrophosphate, 1 mM b-glycerophosphate). The lysates were sonicated at 15W output with 3 bursts of 15 sec each. The protein concentration was estimated according to the BCA Protein Assay Kit (#7780, Cell Signaling Technology). Then the lysate was treated with reducing agent DTT (4.5 mM) and alkylating agent iodoacetamide (IAA) (10 mM). Following this, 1mg protein was digested with PTMScan® Trypsin, TPCK-Treated (#56296, Cell Signaling Technology) overnight at room temperature (RT) on rotation. The digested cell lysate was then purified with C18 reversed-phase Sep-Pak^TM^ columns. Briefly, the digested sample was treated with 1% trifluoroacetic acid (TFA) for acidification (pH should be less than 3) and allowed for precipitation for 15 min on ice. The acidified peptide was centrifuge @ 4.000 x g for 15 min at RT, and the supernatant was collected. The C18 columns were pre-wet with 0.5 ml 100% acetonitrile (ACN) and equilibrated with 1 ml of solvent A (0.1% TFA, 2x). The acidified supernatant was loaded on the column and allowed to pass by gravity. Following this, the column was washed with 1 ml of solvent A (0.1% TFA, 2x) and 0.5 ml of wash buffer (5% ACN/0.1% TFA). The bound peptides were eluted with a sequential wash of 0.2 ml of solvent B (0.1% TFA, 50% ACN) and dried in a vacuum concentrator (Speed-Vac). The dried peptides were then subjected to immunoaffinity purification (IPA). Briefly, approximately 1 mg of dried peptide was resuspended in 1.5 ml of 1X HS IAP Bind Buffer (#25144, Cell Signaling Technology), pH lower than 7.0, and centrifuged @ 10,000 x g for 5 min at 4 °C. The PTMScan® HS Immunoaffinity (K-ε-GG) Magnetic Beads were washed with ice-cold PBS (1X) for 4x. Then the soluble peptide solution was added to the tube containing antibody beads and incubated on an end-over-end rotator for 2h at 4 °C. The tube was centrifuged briefly (2,000 x g, 2-5 sec). The magnetic beads were separated by placing the tube in the magnetic separation rack for 10 sec, and the unbound peptides were discarded. The peptide-bound magnetic beads were washed with ice-cold 1X HS IAP Wash Buffer (#42424, Cell Signaling Technology), and the wash step was repeated for a total of 4x, followed by a wash with ice-cold LCMS-grade water for 2x. The bound peptides from magnetic beads were eluted by adding 50 μl of elution buffer (0.15% TFA) for 10 mins while mixing gently for every 2-3 mins. The eluted sample was removed by placing the tube in a magnetic separation rack. The elution step was repeated, and both eluents were pooled in a single tube. The elute was dried in a vacuum concentrator (Speed-Vac), and the dried peptide was stored at −80°C until LCMS analysis. The LC-MS/MS was performed by the Mass Spectrometry Core in the Research Resources Center of the University of Illinois at Chicago. The dried samples were resuspended in 20 μl 5% CAN, 0.1% formic acid buffer, and 2 μl of the reconstituted sample were analyzed using a Q Exactive HF mass spectrometer coupled with an UltiMate 3000 RSLC nanosystem with a Nanospray Frex Ion Source of protein and peptide with 1 minimum peptide count (Thermo Fisher Scientific). Search results were entered into Scaffold Q+S software (v5.3.4, Proteome Software, Portland, OR) and proteins were identified at a 1% false discovery rate (FDR). The ubiquitylation sites on Akt1 were deduced using scaffold software free viewer version 5.3.3 (http://www.proteomesoftware.com/products/demo-data).

### Statistical analysis

Statistical significance was determined using Student’s *t* test for two-group comparisons or ANOVA followed by Tukey’s or Bonferroni’s post-hoc tests for multiple comparisons. Quantitative data in bar graphs are presented as mean values ± SEM. Results with *p-*values < 0.05 were considered statistically significant. All statistical analyses were conducted using GraphPad Prism 9 software.

## Acknowledgements

This work was funded by the United States NIH grant P01HL160469. The schematic diagrams were made using Biorender.com

## Author contributions

M.O.A: conceived, study design and execution, data analysis and interpretation, and article writing; G.C.H.M, study design and execution, and fund acquisition; L.J. and H.C, performed mass spectrometry and data analysis; A.B.M, study design, article writing, and fund acquisition; C.T. fund acquisition, conceived, study design, data interpretation, supervision, and article writing.

---

**Supplemental Figure 1.**
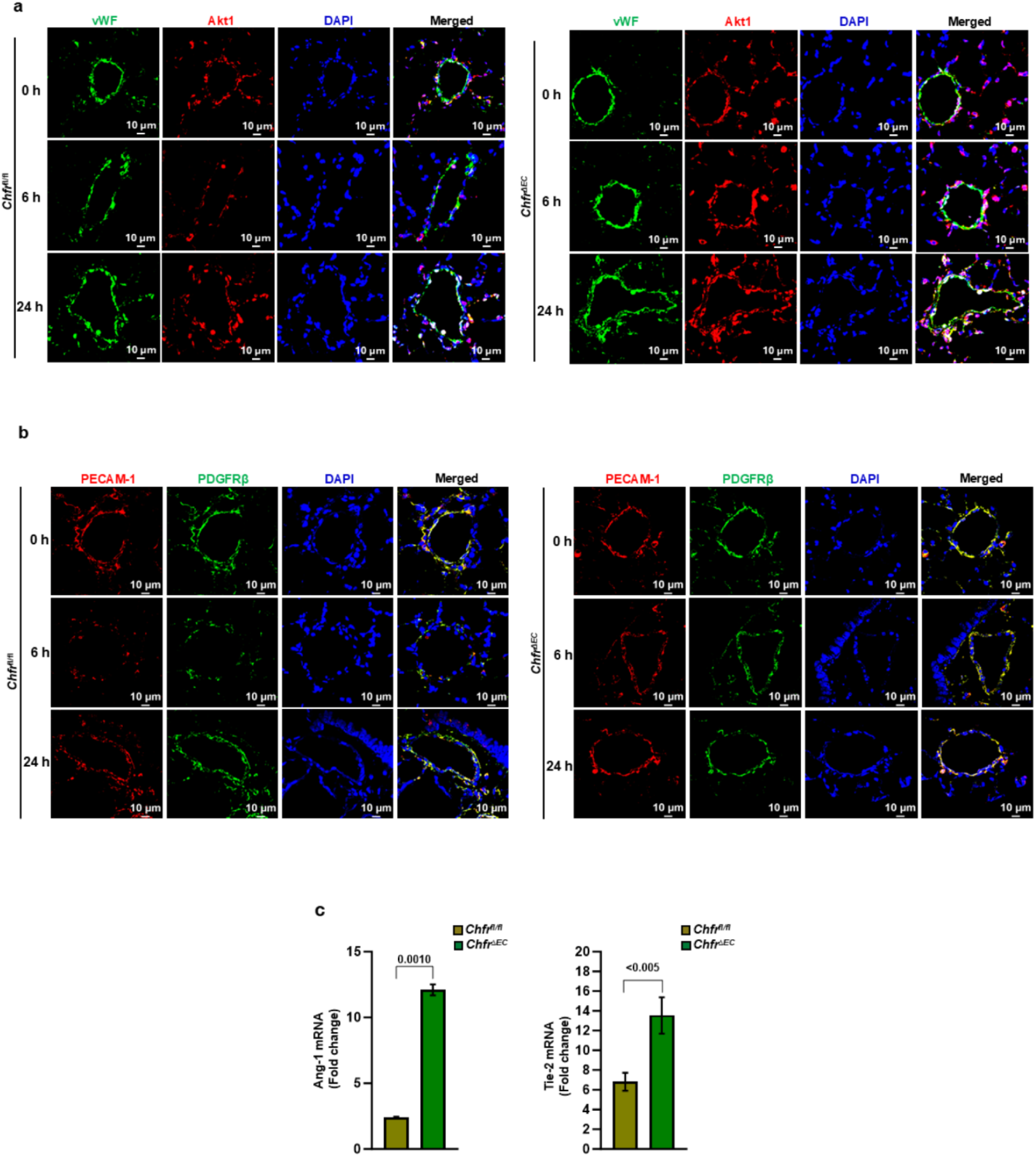
EC-restricted *Chfr* deletion in mice, prevents LPS-induced degradation of Akt1 and pericyte loss in lung vasculature. **a** Lung tissues from *Chfr^fl/fl^* and *Chfr^ΔEC^* mice were collected at 0 and 6 h after i.p. injection of LPS (10 mg/kg bodyweight). Lung sections were stained with antibodies specific to vWF (EC marker, green) and Akt1 (red). **b** Lung tissues from *Chfr^fl/fl^* and *Chfr^ΔEC^* mice were collected at 0 and 6 h after i.p. injection of LPS (10 mg/kg bodyweight). Lung sections were stained with antibodies specific to PECAM1 (EC marker, red) and PDGFRβ (pericyte marker, green). **c** Lung tissues from *Chfr^fl/fl^* and *Chfr^ΔEC^* mice were used for RT-qPCR to measure mRNA expression of Ang1 and Tie-2.

**Supplemental Figure 2.**
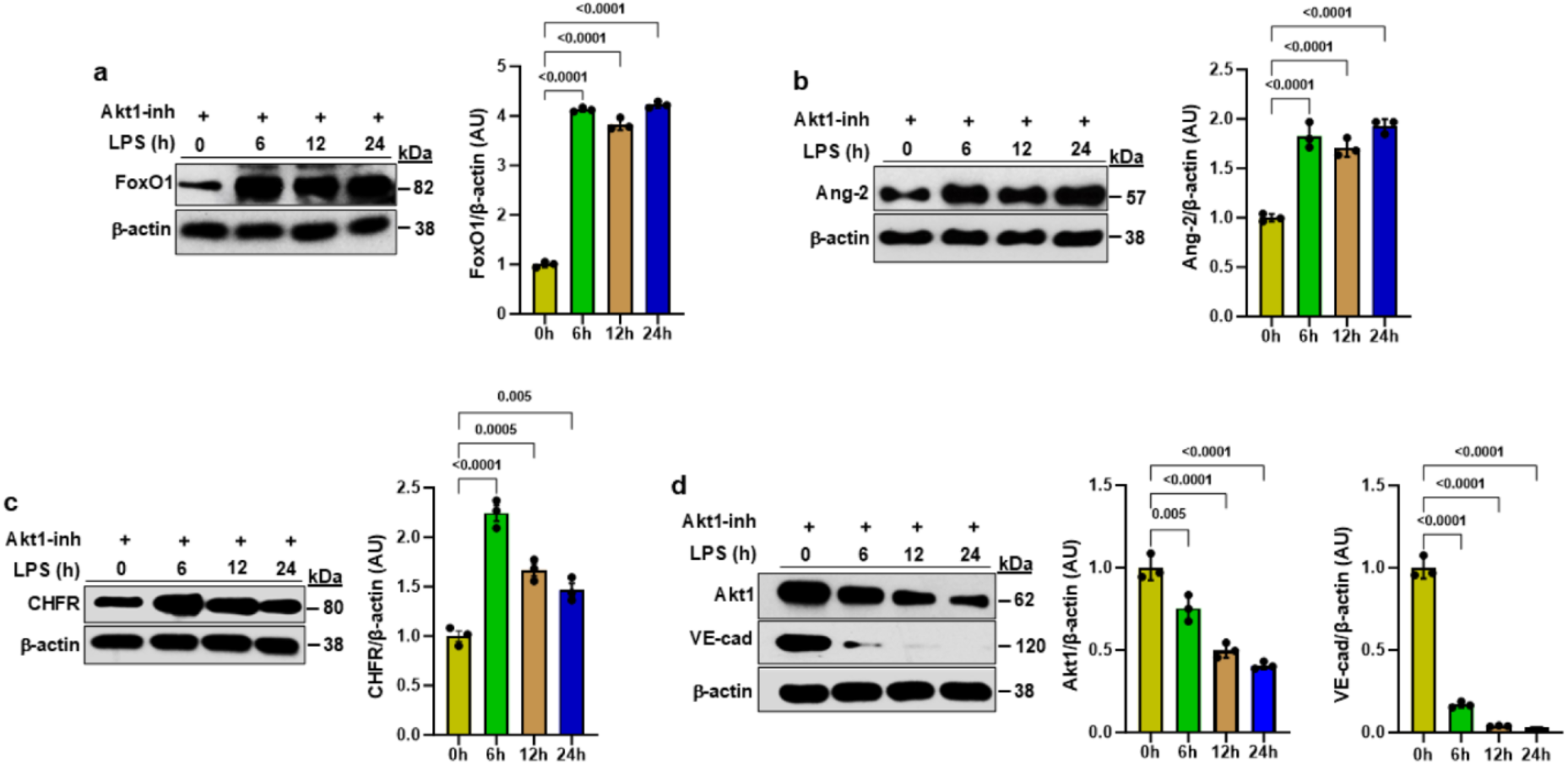
Pharmacological inhibition of Akt1 augments LPS-induced expression FoxO1, Ang-2, and CHFR, and suppresses the expression of Akt1 and VE-cadherin in EC. **a-d** HLMVEC were pretreated with the Akt1 inhibitor (Akt1-inh) for 1 h before stimulation with LPS (5 μg/ml) for 0, 6, 12, and 24 h. IB analysis showed increased expression of FoxO1, Ang-2, and CHFR, whereas expression of Akt1 and VE-cadherin was suppressed. Shown are mean values ± SEM (*n* = 3 independent experiments; two-way ANOVA followed by Tukey’s post hoc test).

**Supplemental Figure 3.**
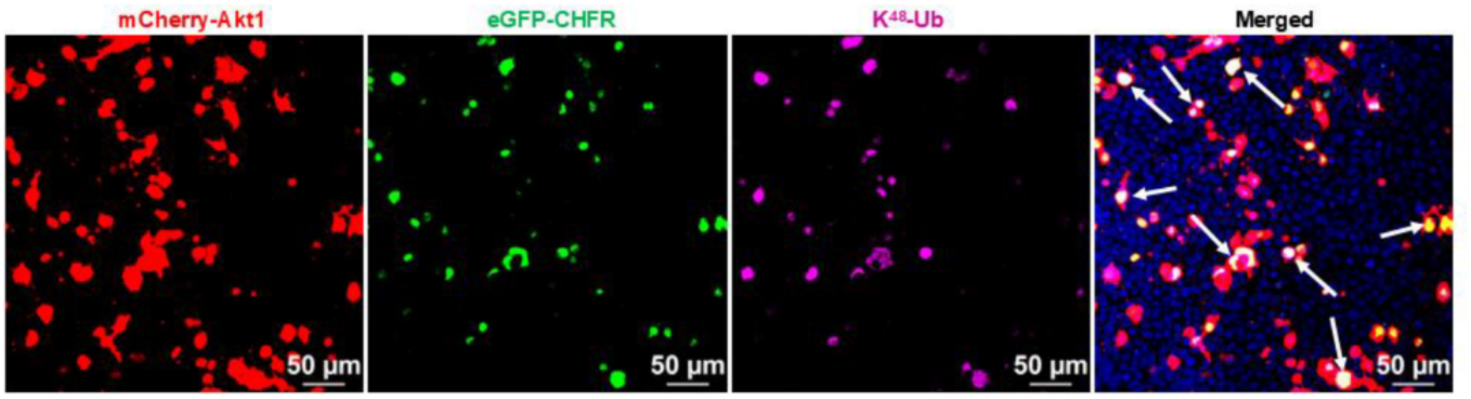
I*n vitro* validation of ubiquitylation of Akt1 by E3 ligase CHFR: HEK-293T cells were transfected with pmCherry-Akt1 (WT), eGFP-CHFR (WT), and HA-Ub (WT). The transfected cells were stained with K^48^-linkage specific antibody. Confocal imaging showing the co-localization of HA-Ub (WT), pmCherry-Akt1 (WT), and eGFP-CHFR (WT).

**Supplemental Figure 4.**
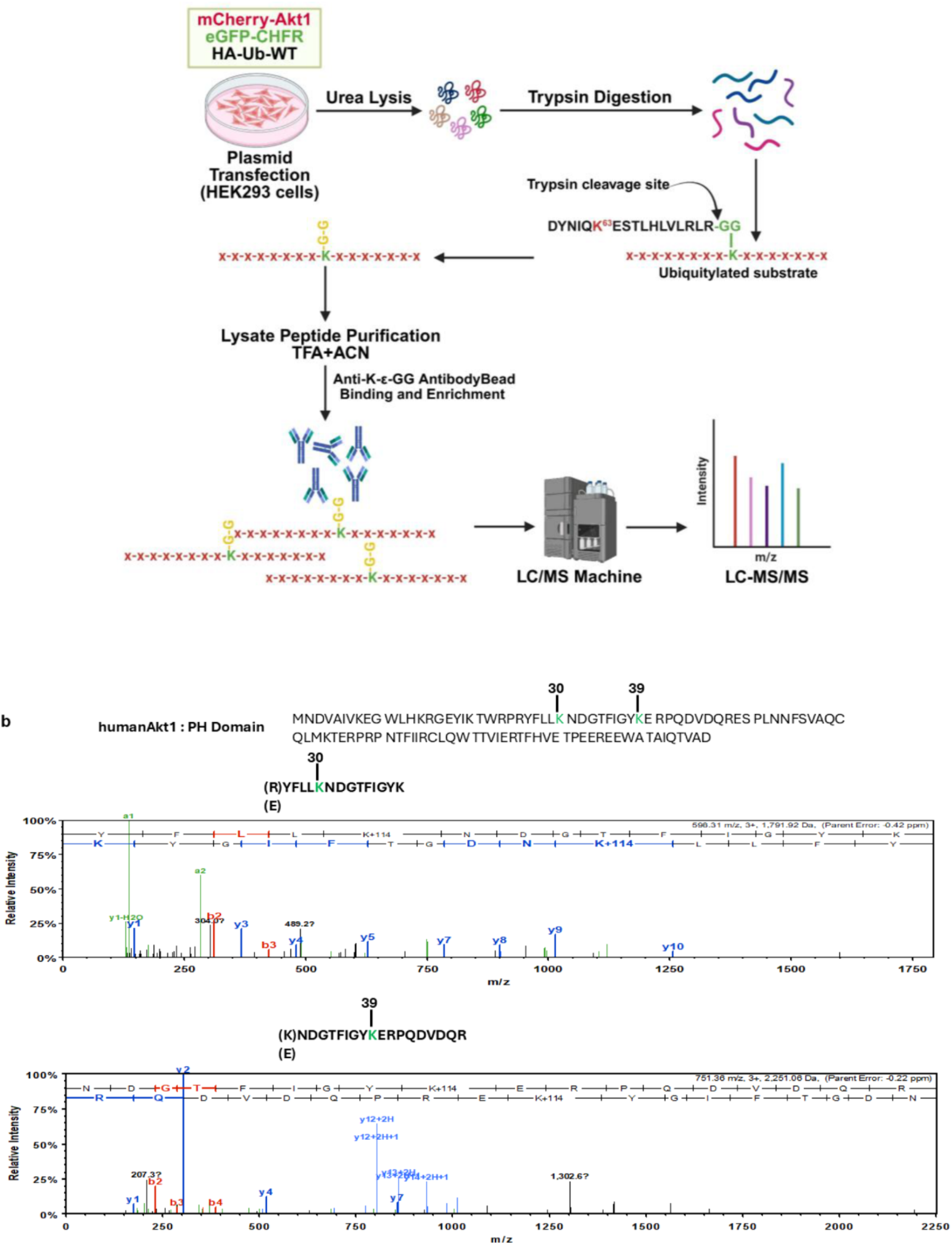
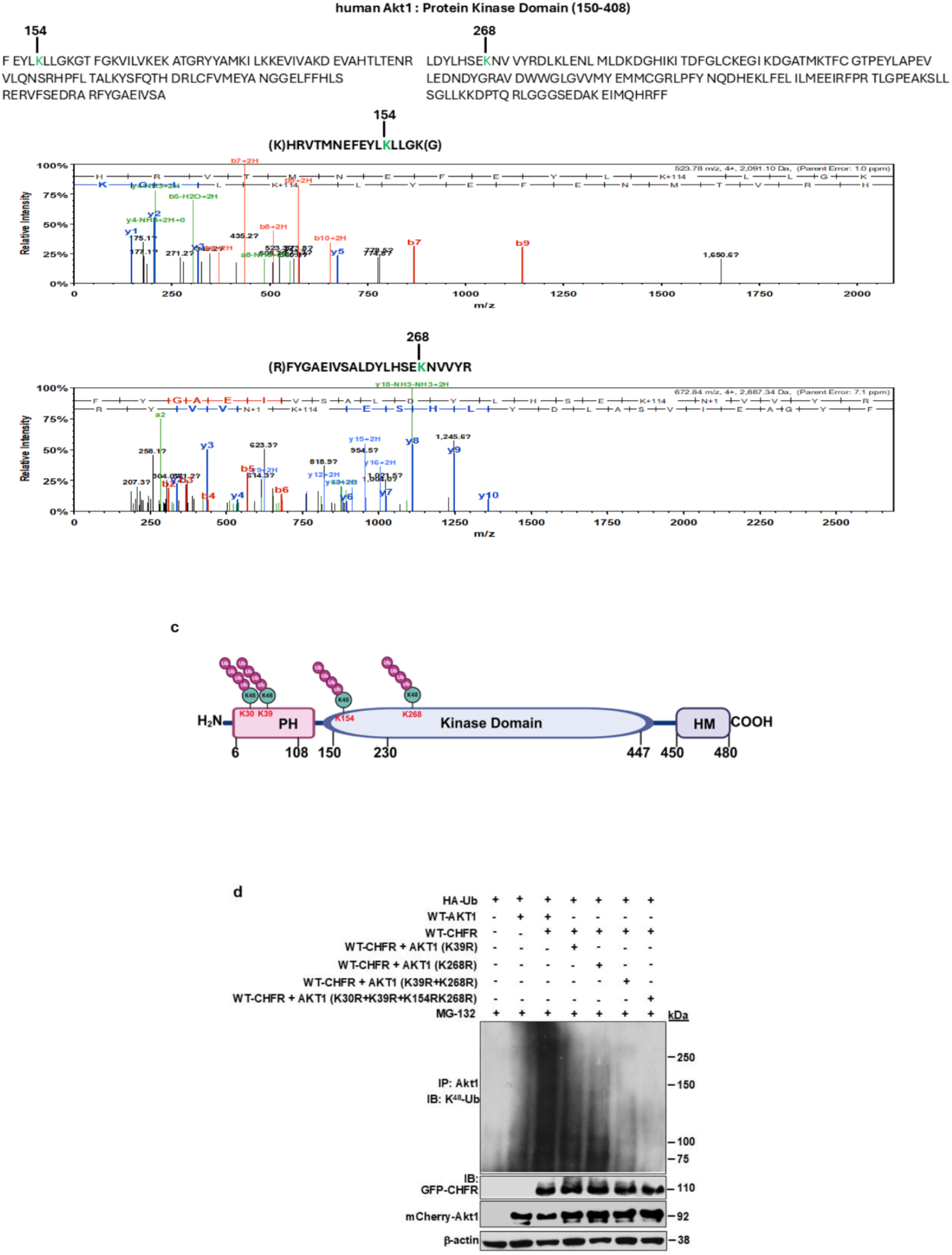
Identification of ubiquitylation sites on Akt1. **a** schematic showing the workflow for LC-MS analysis. HEK293 cells were transfected with pmCherry-Akt1 (WT), eGFP-CHFR (WT), and HA-Ub (WT). At 48 h post-transfection, cells were collected and subjected to urea lysis followed by trypsin digestion. The digested peptides were affinity-purified and immunoprecipitated with anti-di-glycine remnant antibodies to enrich for ubiquitylated peptides, followed by LC-MS/MS analysis. **b** shows the Akt1 ubiquitylated “K” containing peptides identified by LC-MS. **c** Model of Akt1 protein structure showing ubiquitylation sites. **d** HEK-293T cells transfected with WT-HA-Ub (0.5 μg/ml) alone or co-transfected with WT-CHFR (1.5 μg/ml), WT-Akt1 (1.5 μg/ml), or Ub-defective Akt1 (K30R, K268R, K39R+K268R, and K30R+K39R+K154R+268R) plasmids were used to study ubiquitylation of Akt1. IB analysis showed that the CHFR-induced ubiquitylation of Akt1 was blocked in “K to R” Akt1 mutants expressing cells compared with WT-Akt1.

